# A Synthetic Peptide Encoded by a Random DNA Sequence Inhibits Discrete Red Light Responses

**DOI:** 10.1101/669564

**Authors:** Tautvydas Shuipys, Raquel F. Carvalho, Maureen A. Clancy, Zhilong Bao, Kevin M. Folta

## Abstract

We have identified a synthetic peptide that interrupts discrete aspects of seedling development under red light. Previous reports have demonstrated that plants transformed with random DNA sequences produce synthetic peptides that affect plant biology. In this report one specific peptide is characterized that inhibits discrete aspects of red-light-mediated *Arabidopsis thaliana* development during photomorphogenesis. Seedlings expressing the PEP6-32 peptide presented longer hypocotyls and diminished cotyledon expansion when grown under red light. Other red-light-mediated seedling processes such as induction of *Lhcb* (*cab*) transcripts or loss of vertical growth remained unaffected. Long-term responses to red light in PEP6-32 expressing plants, such as repression of flowering time, did not show defects in red light signaling or integration. A synthesized peptide applied exogenously induced the long-hypocotyl phenotype under red light in non-transformed seedlings. The results indicate that the PEP6-32 peptide causes discrete cell expansion defects during early seedling development in red light, mimicking weak *phyB* alleles in some aspects of seedling photomorphogenesis. The findings demonstrate that new chemistries derived from random peptide expression can modulate specific facets of plant growth and development.

**One Sentence Summary:** A plant line expressing random DNA sequence expresses a synthetic peptide that affects specific red-light responses in a developing seedling.

## Introduction

There is significant interest in identifying new molecules that modulate plant growth and development, as well as protect them from biotic and abiotic stress. Efforts in chemical genomics have identified novel compounds with specific interactions in the plant, including flowering regulators (Fiers et al., 2017), exocytosis inhibitors (Zhang et al., 2016), and activators of hormonal signaling (De Rybel et al., 2009). Synthetic molecules that serendipitously excite specific biological processes provide a means to create phenotypes in plants that bypass barriers of genetic redundancy or lethality (Hagihara et al., 2019). These approaches drive discovery of new connections between chemistry and biology with the goal of devising new strategies for plant protection, agricultural productivity, or directed synthesis of particular secondary metabolites.

Another approach sought to identify novel regulators in populations of transgenic plants where each plant expressed a unique peptide encoded by randomized DNA information (Bao et al., 2017). *Arabidopsis thaliana* plants were transformed with libraries of constructs bearing the random DNA sequence, flanked by start and stop codons, and driven by the constitutive CaMV35S promoter. Populations of transformed plants presented a surprisingly frequent number of reproducible phenotypes, apparently due to expression of the random DNA sequence, or potentially its encoding RNA. The results of Bao et al., (2017) identified a new candidate peptide that showed conditional lethality, and another (noted as PEP6-32) that affected seedling growth in red light, but not under blue or far-red wavelengths.

Red light is sensed through the light sensor family known as the phytochromes (reviewed in Quail, 2002; Xu et al., 2015). The phytochrome B sensor (phyB) has a central role in red light responses (Reed et al., 1993; Parks and Spalding, 1999). In the developing seedling phyB effects hypocotyl elongation, cotyledon expansion, orientation to the gravitational vector, chloroplast development and gene expression (Somers et al., 1991; Whitelam et al., 1998; Tepperman et al., 2004; Chen and Chory, 2012). Later in development phyB is central in response to shaded environments (Whitelam and Smith, 1991; Robson et al., 1993) and has an important role in the transition from vegetative growth to flowering (Valverde et al., 2004).

The present work attempts to describe where the PEP6-32 peptide interacts with plant biology. The central question is if the peptide is interfering with phyB signaling directly or if it is interfering with an aspect of seedling development only observed under red light illumination. Arabidopsis has been extensively characterized for early light responses, and hosts a great array of genetic tools. Therefore it is possible to test the hypothesis that the PEP6-32 peptide is specifically interfering with phyB-regulated light signaling by characterizing a suite of phyB-related responses.

This report presents a comprehensive physiological characterization of the effect of the PEP6-32 peptide on red light signaling and response. Such careful characterization is necessary before there can be meaningful efforts to identify the precise mechanism of light-signal attenuation, as identifying interacting molecules requires understanding precisely where and when the interaction is taking place. The work also tests the possibility of using this synthetic sequence as an exogenous growth regulator in interfering with red light response.

## Results

### *PEP6-32* expressing seedlings exhibit elongated hypocotyls under red light

The initial characterization of the *PEP6-32* line revealed that the seedlings exhibited decreased hypocotyl growth inhibition under red light, but not under far-red or dark-growth conditions (Bao et al, 2017). Furthermore, there was only a slight difference in hypocotyl length under low fluence blue light (≤0.5 μmol⋅m^−2^⋅s^−1^) that was absent under higher fluence rates. These results suggested that the peptide might be interfering with light sensing or response through phytochrome B (Neff and Chory, 1998). To further explore this possibility, transgenic lines were compared to non-transformed controls and *phyB-5* mutant seedlings. These *phyB* mutants contain substitution W552STOP in the coding sequence resulting in a non-functional, truncated phyB molecule (Reed et al., 1993), crossed against a Col-0 *phyA* mutant, and identified in subsequent generations for sensitivity to red light after backcrosses to Col-0 (as described in Wang et al, 2013) providing an important null-allele control for comparison.

Non-transformed controls, several independent transformants expressing the *PEP6-32* peptide, and *phyB* were grown vertically in light chambers under constant red light (30 μmol⋅m^−2^ ⋅s^−1^), constant blue light (10 μmol⋅m^−2^ ⋅s^−1^), or complete darkness for 96 h. Under red light, the peptide expressing plants showed significantly less hypocotyl growth inhibition compared to non-transformed controls (Fig. 1A). However, the difference in relative hypocotyl length was not as great as that observed in *phyB* seedlings. This finding indicates that the peptide does not completely eliminate phyB response (as in the *phyB* mutant) yet it exerted enough influence to significantly alter normal response. Under blue light (Fig. 1B), there was no significant difference between control and peptide expressing plants, while *phyB* did have a slight, but significant increase in relative hypocotyl length, consistent with previous observations (Neff and Chory,1998).

**Figure 1.**
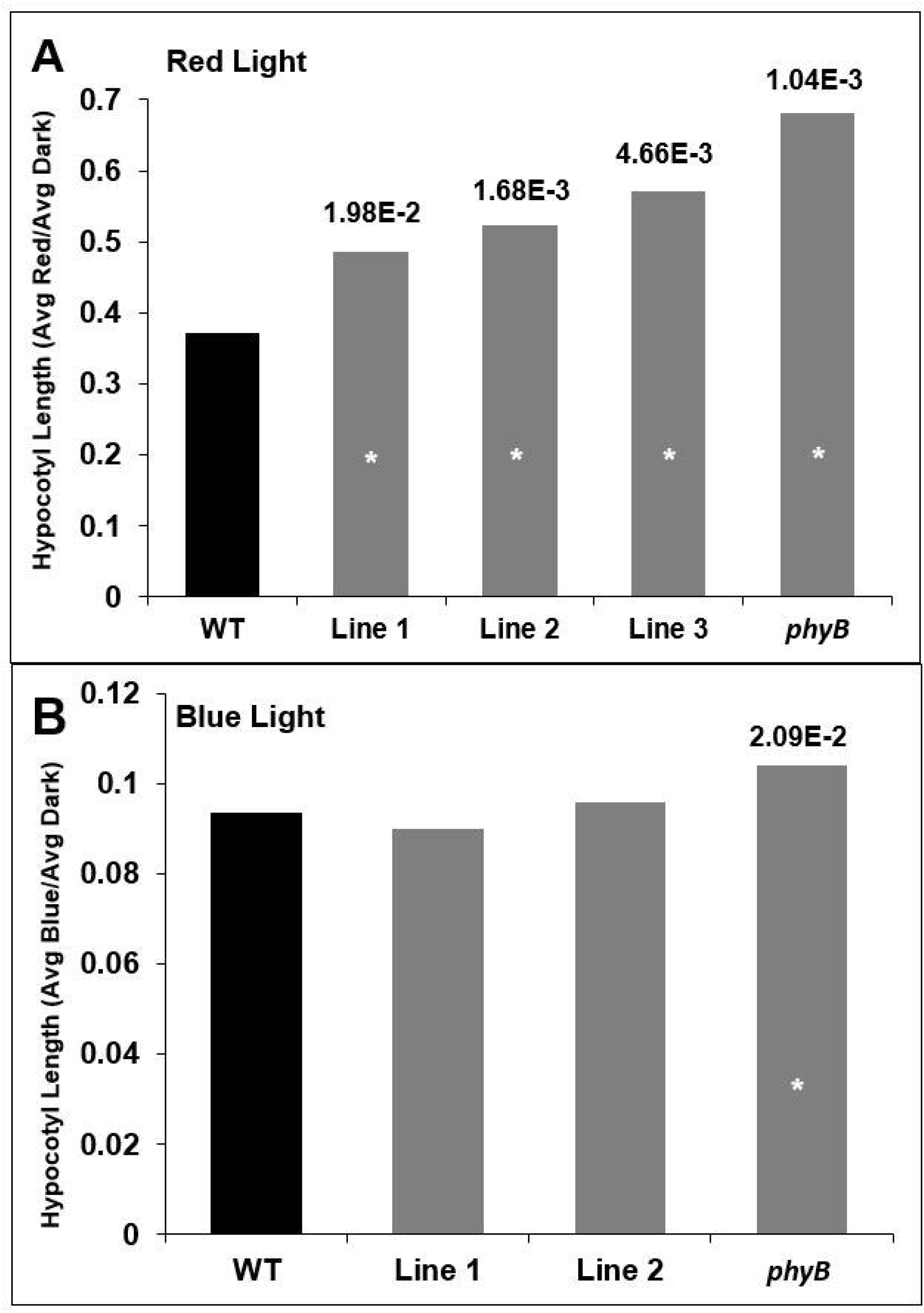
Expression of the *PEP6-32* peptide results in plants with longer hypocotyls than WT but shorter than *phyB* mutants under red light. Relative hypocotyl length of non-transformed control Col-0 (WT), independent transformed lines expressing the PEP6-32 peptide (Lines 1-3), and *phyB* plants under 30 μmol⋅m^−2^⋅s^−1^ red light (A) or 10 μmol⋅m^−2^⋅s^−1^ blue light (B) after 96 h. Relative hypocotyl length is calculated as the average hypocotyl length of light grown plants divided by the average hypocotyl length of dark grown plants. N=9-10 for Lines 1-3, N=16 for *phyB*, and N=17 for WT. Asterisk indicates statistically significant difference between transgenic seedlings compared to wild-type seedlings using the Mann-Whitney U Test; p values are listed above the bars for the significant results.

### Growth against a gravitational vector in red light

In addition to regulating hypocotyl elongation under red light, phyB also acts to guide directional growth of seedlings against the gravitational vector (Poppe et al., 1996). Under constant red light, seedlings grow in various directions, not vertically. This phenotype is not observed in *phyB* seedlings, as they grow straight under red light. This has been shown to be caused by phyB’s inhibition of PIFs which are involved in the development of the gravity sensing endodermal amyloplasts (Kim et al., 2011), and requires participation specific 14-3-3 proteins (Mayfield et al., 2007). To test the response in the PEP6-32 expressing plants, seedlings were grown for 96 h length under constant red light (30 μmol⋅m^−2^ ⋅s^−1^) and compared to *phyB* and wild-type seedlings (Figure 2). For this experiment, the growth direction was measured from 0-180° with 0° being straight up and 180° being directly down, in line with the direction of gravity. As expected, the *phyB* mutants generally grew upright under constant red light with the average deviation of 39.3° from vertical. There was no significant difference in the angle for the peptide expressing plants when compared to WT controls.

**Figure 2.**
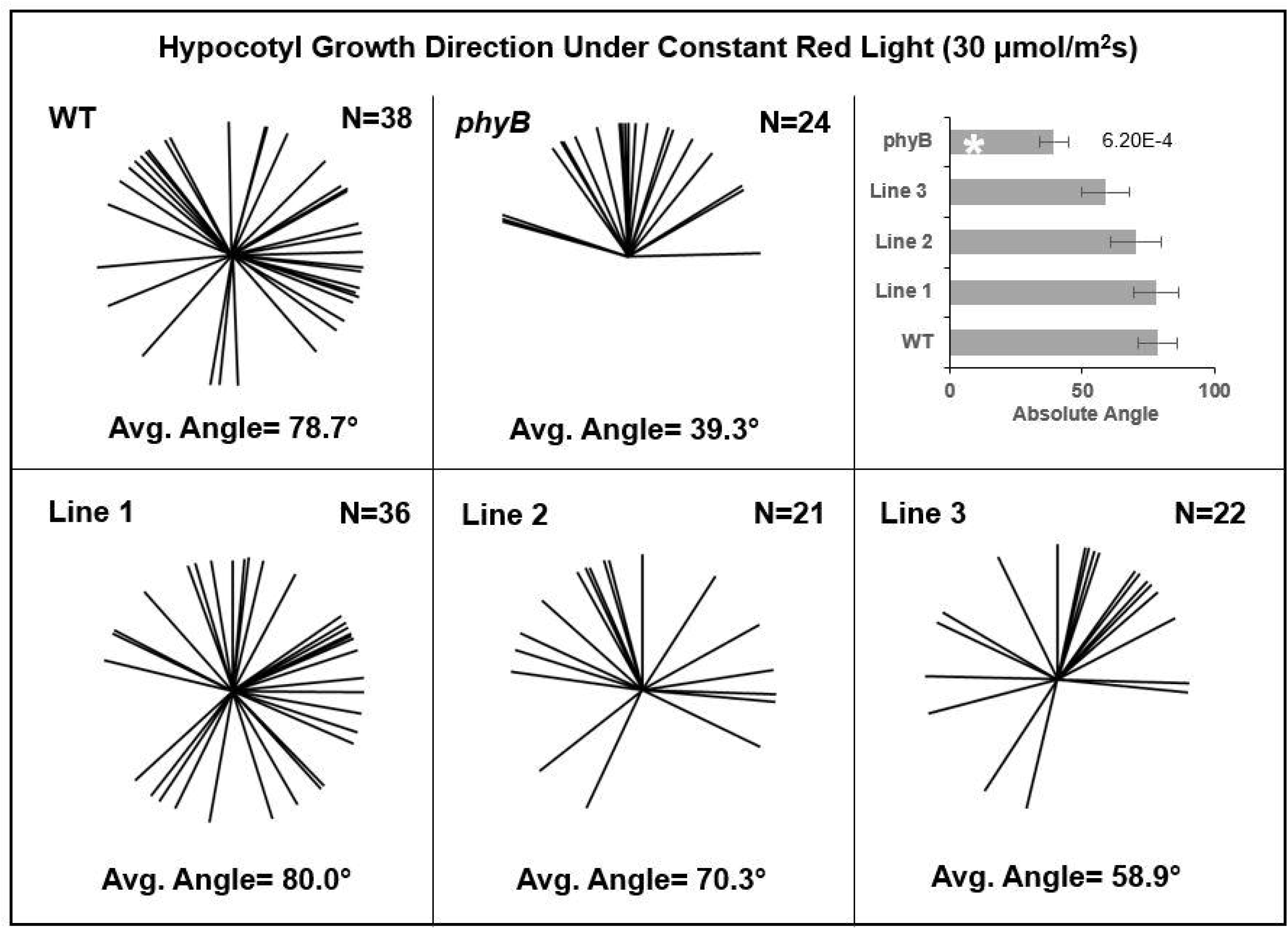
PEP6-32 expressing plants exhibit normal directional growth under red light. Graphical depiction of non-transformed control Col-0 (WT), independent transformed lines expressing the PEP6-32 peptide (Lines 1-3), and *phyB* plants after 96 h of 30 μmol⋅m^−2^⋅s^−1^ red light. Measurements taken from three independent experimental replicates. Sample size for each line is listed in the top right of each box. The average angle of growth is listed at the bottom of each box and is defined as the absolute angle from 0-180 where 0 is straight up (opposite the direction of gravity) and 180 is straight down (in the direction of gravity). Asterisk indicates statistically significant difference between transgenic seedlings and wild-type seedlings using the Mann-Whitney U Test. P values are listed to the right of the bars for the significant results. Error bars indicate standard error of the mean.

### Effect of PEP6-32 on cotyledon expansion

Cotyledon expansion is also observed during photomorphogenic development in response to red light. This process has been shown to be controlled redundantly by both phytochromes (phyA and phyB) and cryptochrome (cry1) with phyB and cry1 the major components under white light, phyB under red light, and cry1 under blue light (Neff and Chory, 1998). Seedlings expressing *PEP6-32* were grown under 100 μmol⋅m^−2^⋅s^−1^ white light, 30 μmol⋅m^−2^⋅s^−1^ red, or 10 μmol⋅m^−2^⋅s^−1^ blue light with a 16 h photoperiod for 7 d. Afterwards, the cotyledons were removed from the plants and imaged. The surface area was calculated using ImageJ. The results in Figure 3 show that under white or blue light *phyB* seedling cotyledons were smaller than wild type, yet the *PEP6-32* lines were the same size or slightly larger. Under red conditions the *PEP6-32* lines showed a significantly decreased area, but not to the same extent as *phyB*. Interestingly, the cotyledon size of the two independent *PEP6-32* lines was slightly, but significantly greater than non-transformed controls under blue light, in contrast to *phyB* plants that are smaller than wild-type seedlings. The *PEP6-32* Line 2 genotype did not show significant differences from wild type under any conditions.

**Figure 3.**
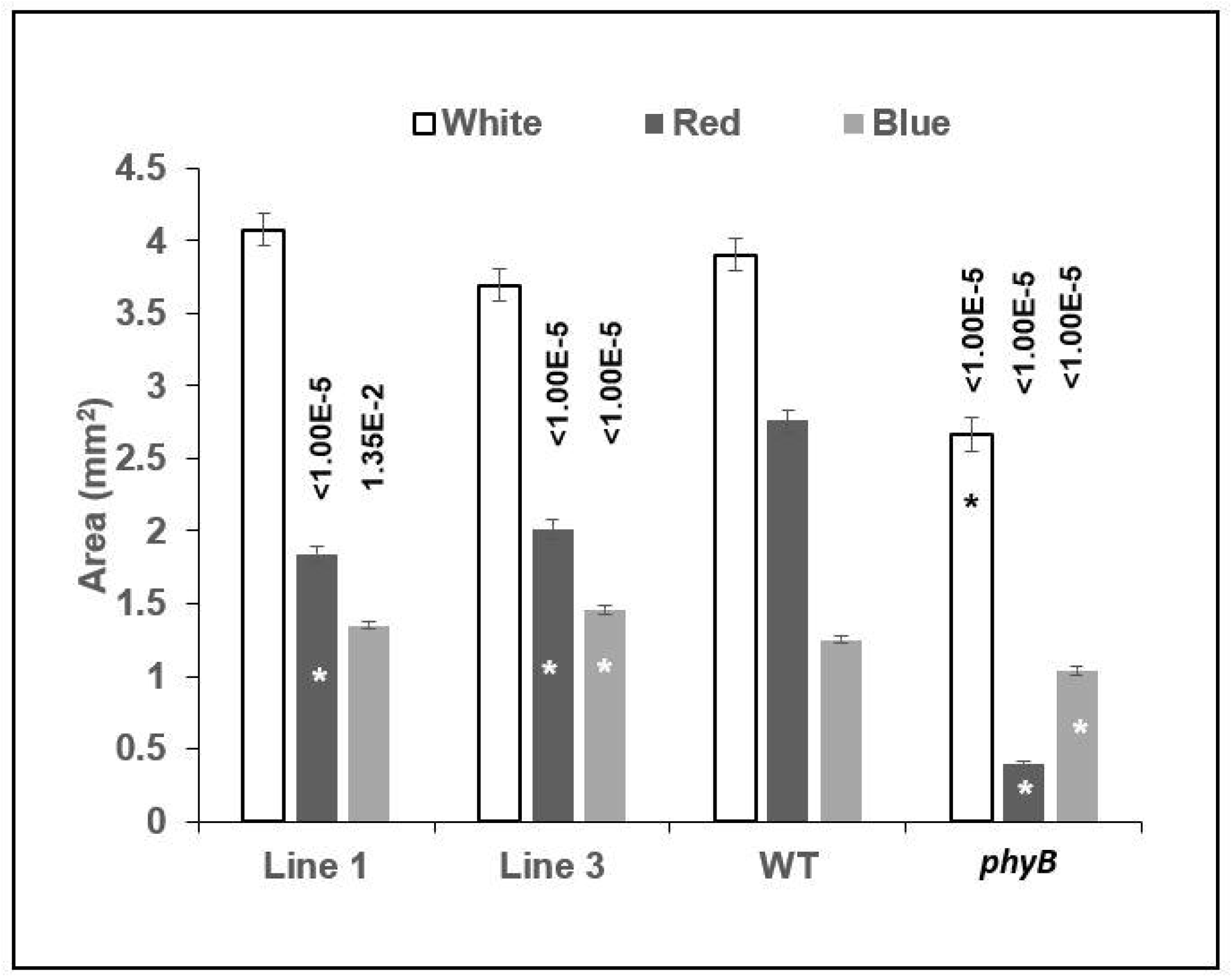
PEP6-32 expressing plants exhibit smaller cotyledons under red light. Cotyledon size of non-transformed control (Col-0), independent transformed lines expressing the PEP6-32 peptide (Lines 1 and 3), and *phyB* plants after 7 d under long day conditions. The fluence rate of white light was 96 μmol⋅m^−2^⋅s^−1^, red light was 30 μmol⋅m^−2^⋅s^−1^, and blue light was 10 μmol⋅m^−2^⋅s^−1^. The results shown here are an average of two independent experiments, N=73-82 per line. ImageJ was used to determine area of each cotyledon in mm^2^. Asterisks indicate statistically significant difference between transgenic or mutant seedlings compared to wild-type seedlings using the Mann-Whitney U Test. P values are listed above the bars for the significant results. Error bars indicate standard error.

### Differences in red light regulated gene expression

If the peptide interacts with phyB photosensor chemistry or signal transduction directly, changes in the timing or amplitude of phyB-mediated transcript accumulation would be observed. Transcript accumulation corresponding to two red-light regulated members of the *Light Harvesting, Chlorophyll-Binding* (*Lhcb* or *Chlorophyll a/b Binding;* cab) gene family (*CAB2* and *LHCB1*5)* was monitored in response to a short red light treatment. Four-day old etiolated seedlings were treated with 8 μmol⋅m^−2^⋅s^−1^ of red light for 1 h or kept in complete darkness as a control. The results show that there was no significant difference in accumulation of either of the two transcripts in the *phyB* mutant seedlings (Fig. 4D), while wild-type seedlings exhibited a typical and significant induction of both (Fig. 4C). The *PEP6-32* seedlings showed normal induction of both transcripts (Fig. 4A,B) comparable to wild-type, non-transformed seedlings.

**Figure 4.**
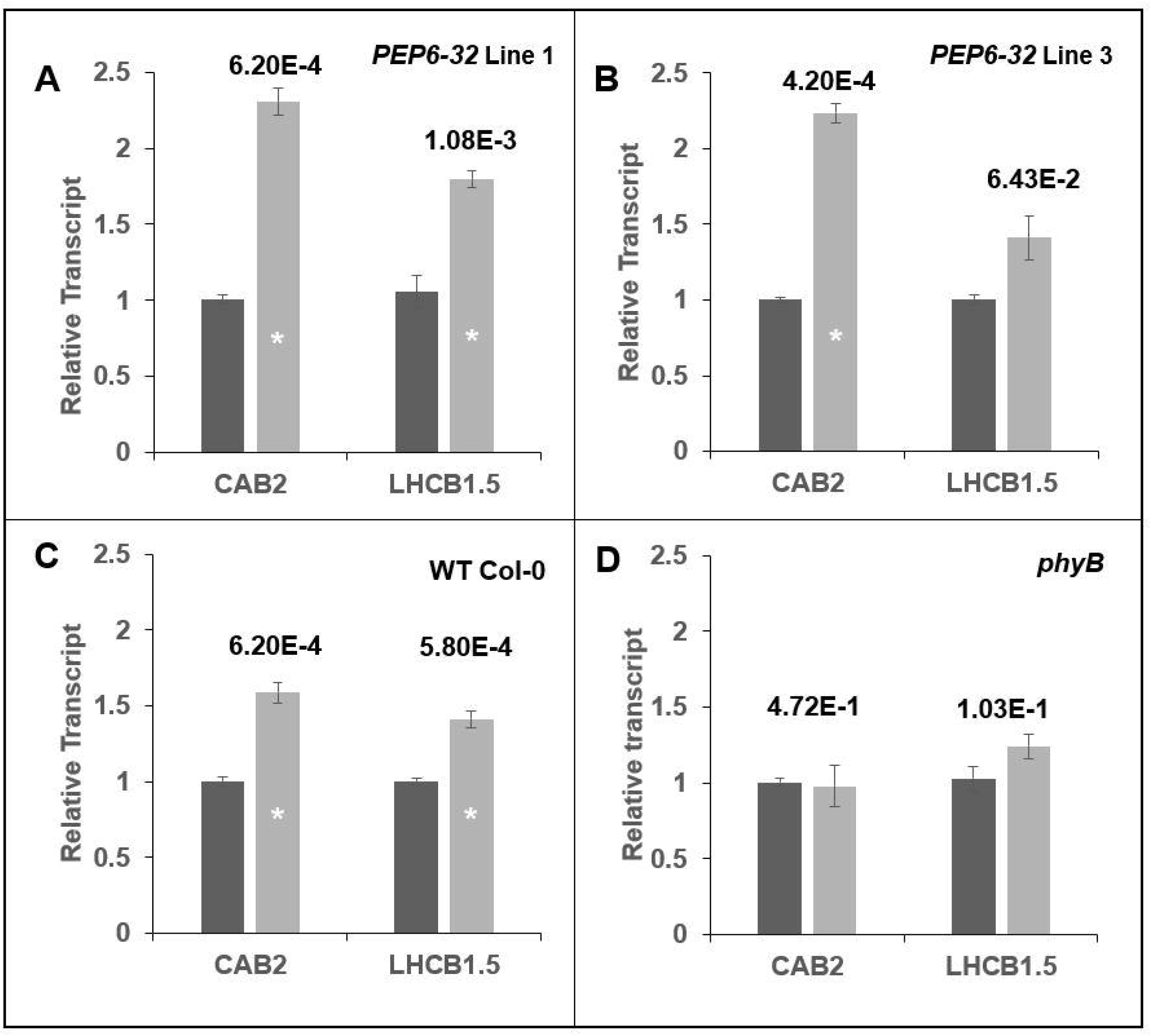
Expression of the peptide does not affect induction of *CAB2* or *Lhcb1*5*. Fold change in the steady-state accumulation of *CAB2* and *Lhcb1*5* transcripts in independent peptide expressing lines (A,B), compared to a non-transformed control (Col-0; C), and *phyB* mutant seedlings (D). Transcript levels were measured using RT-qPCR after a 1 h 8 μmol⋅m^−2^⋅s^−1^ red light treatment of 4 d old etiolated seedlings relative to dark-grown controls. Results shown here are an average of three independent experimental replicates and three technical RT-qPCR replicates. The *AtUFP* transcript was used as a constitutively expressed reference. Error bars indicate standard error of the mean. The asterisk indicates a statistically significant difference between transgenic and non-transformed controls Mann-Whitney U Test. P values are listed above the bars for the significant results.

### Peptide effect on the flowering time

The phyB photoreceptor also has a central role in the regulation of flowering. To test if the PEP6-32 peptide affected phyB (or other) response during the floral transition Col-0, *phyA* (far-red sensing impaired)*, phyB*, and segregating positive peptide and non-transgenic seedlings were transplanted into pots 9 d after germination. The plants were grown in a growth chamber under long day conditions, and then flowering was scored when the inflorescence was 1 cm in length or greater. The results show that instead of trending towards early flowering, the PEP6-32 lines trend towards slightly later flowering (Fig. 5A). The *phyB* mutant plants flowered about 4 d before Col-0, but the peptide lines flowered 2-4 d later. Two PEP6-32 expressing lines exhibited a short, but significantly later time to flower while two others did not. The corresponding mRNA levels were examined and indicate that steady-state transcript levels did not correlate with the slight delays observed in the timing of flowering (Fig. 5B)

**Figure 5.**
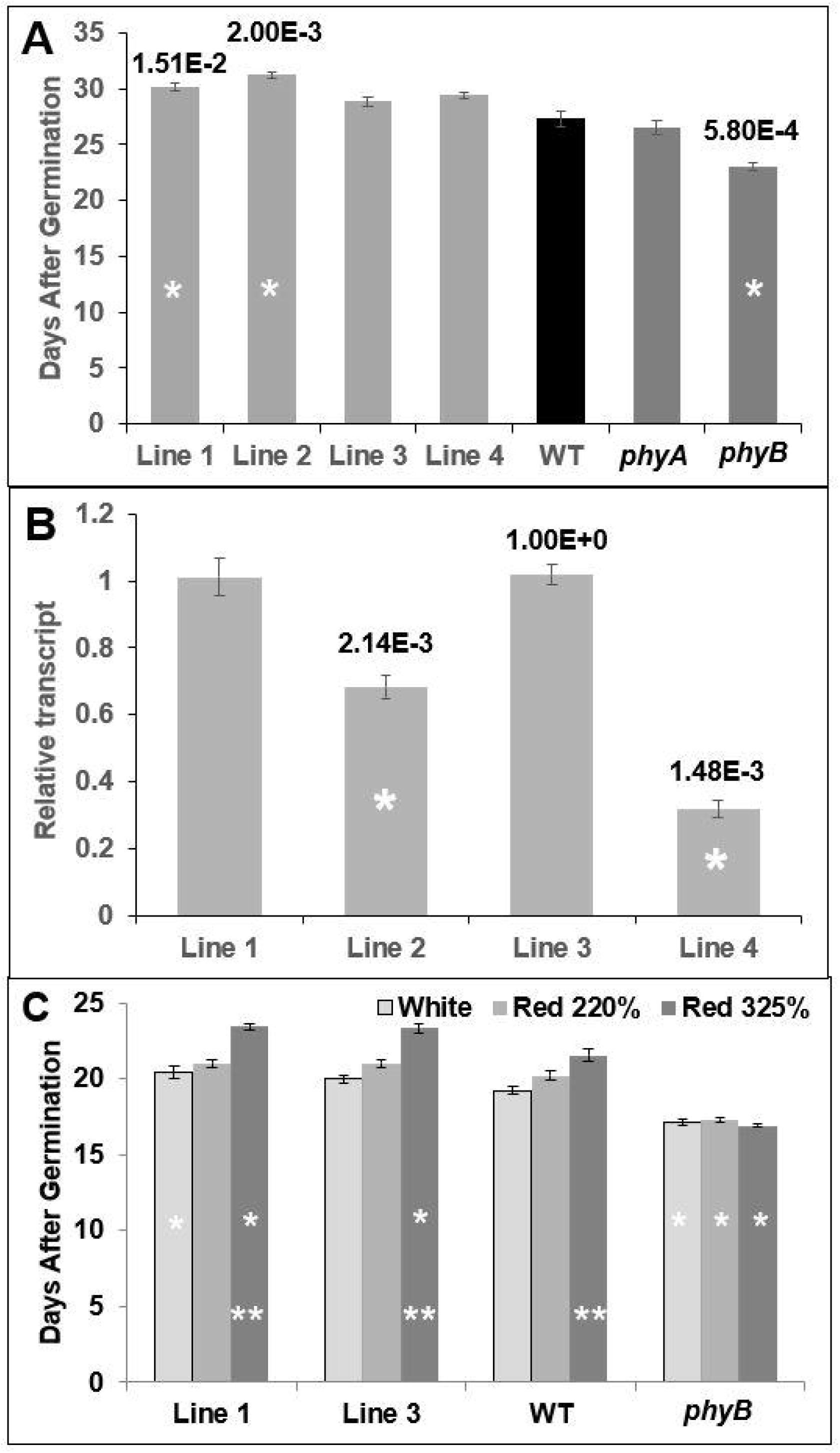
The peptide does not induce early flowering. (A) Examining the time to flower of non-transformed control (Col-0), independent transformed lines expressing the PEP6-32 peptide (Lines 1-4), *phyA* plants, and *phyB* plants in a growth chamber at 21°C under long day conditions. The data indicate the average number of days until flowering. The asterisk indicates a statistically significant difference from wild-type plants. P values are listed above the bars for all significant results. (B) Levels of PEP6-32 transcript in the different independent transformed lines relative to Line 1, as measured using RT-qPCR. Results shown here are an average of two independent RT-qPCR experiments comprised of three technical replicates. The asterisk indicates a statistically significant difference compared to Line 1. All p values are listed above the bars. (C) The effect of increasing proportion of red light on flowering time. Lines 1, 3, WT, and *phyB* were grown in LED light chambers set to three conditions designed to mimic the light spectrum of the growth chamber. In the treatments the proportion red light was increased to 220%, or 325% of normal (see Supplemental Fig. 1). A single asterisk indicates statistically significant difference compared to wild-type seedlings. The double asterisk indicates a statically significant difference from plants grown under normal white light for each respective line. P ≤ 0.05. Error bars indicate standard error of the mean.

Another way to examine the potential role of PEP6-32 in phyB-mediated signalling during flowering would be to monitor flowering time in relevant genotypes in red-enriched light environments. Increasing the amount of red light should amount to increased photoactivated phyB and result in delayed flowering. For this experiment plants were grown under LED light containing blue, green, amber and red light (450, 530, 590 and 660 nm, respectively) and the same conditions with an increased the amount of red light (660nm) at the cost of amber light (590nm) to maintain constant photosynthetically active radiation (Supplemental Fig. 1). Three sets of treatments were used, red, blue, green, amber at 100 µmolm^−2^s^−1^, the same treatment with 220% red, and the third treatment with 325% red. Under all three conditions, *phyB* mutant plants showed no change in the time to flower as expected (Fig. 5C). At 325% red, both peptide expressing lines showed an increased time to flower which was significantly different from non-transformed controls as well as different from the control white light grown plants. Increasing red to 220% had a slight, but not significant difference in time to flower for both WT and peptide expressing lines as compared to the control white light treatment. These data indicate that the peptide is not impairing phyB activity with respect to flowering time (Fig. 5A,C).

### Exogenous application of the peptide to non-transformed plants

It was important to test if a synthetic growth regulator like PEP6-32 could modulate its effects when applied externally. To test this, a synthesized version of the peptide was added to solid and liquid media at relatively high concentrations and hypocotyl growth inhibition was measured after exposure to constant red light illumination. Fluence rate/response tests were performed to test the effect of exogenously applied peptide under these conditions.

The synthesized peptide was added to buffer at 1 μM in solid minimal media. The seedlings were grown vertically under 1, 5, 10, 30, or 100 μmol⋅m^−2^⋅s^−1^ of constant red light for 96 h. The results reveal that plants grown in the presence of PEP6-32 exhibited in a slight yet significant difference in hypocotyl length at fluence rates of 5 μmol⋅m^−2^⋅s^−1^ or greater (Fig. 6). No significant difference was observed in dark grown plants or those grown under 1 μmol⋅m^−2^⋅s^−1^ red light.

**Figure 6.**
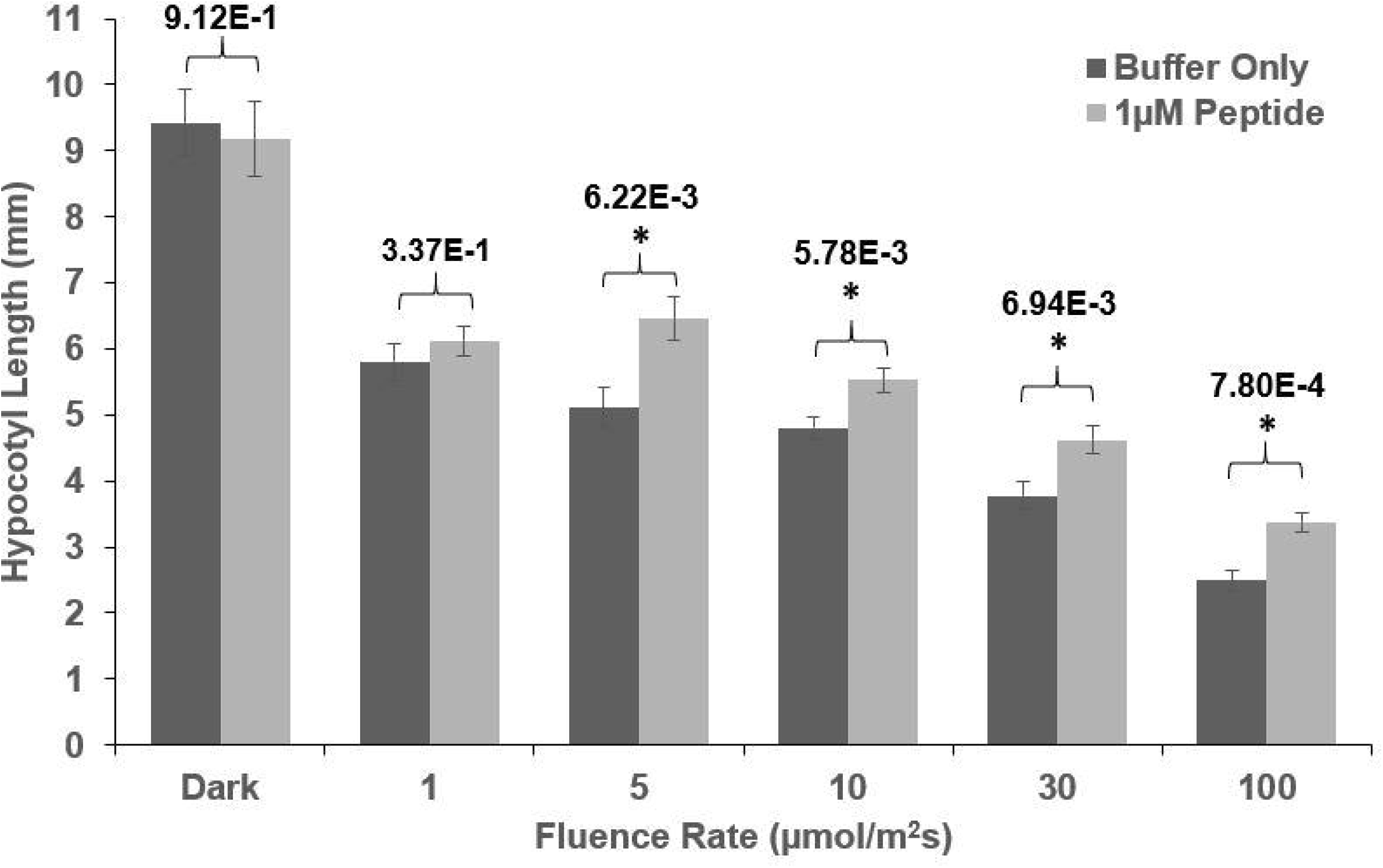
WT plants grown on media supplemented with synthesized PEP6-32 exhibit a slight but significant increase in hypocotyl lengths under constant red light. Average hypocotyl length of non-transformed seedlings grown on minimal media containing 1 μM of peptide (light bars) or a buffer control (dark bars) after 96 h of 1, 5, 10, 30, or 100 μmol⋅m^−2^⋅s^−1^ red light. Data were pooled from multiple independent experiments, N=19-64. The asterisk indicates a statistically significant difference between plants grown in the presences of the PEP6-32 peptide compared to media without the peptide, using Mann-Whitney U Test. All p values are listed above the bars. Error bars indicate standard error of the mean.

## Discussion

The 2017 report by Bao and colleagues provided evidence that new synthetic regulators of plant growth and development could be identified within populations of transgenic plants expressing random DNA sequences. Hundreds of reproducible phenotypes have been observed in plants from these populations, and it now the challenge is to identify the mechanisms by which the peptides (or possibly their nascent RNAs) exert their effects on specific aspects of plant biology.

In the developing seedling, the effects of PEP6-32 mimic those of a weak *phyB* mutant. Hypocotyl growth inhibition and cotyledon expansion are impaired seedlings expressing this peptide. These seedling attributes are observed after 96 h of constant illumination and are consistent with a role for the peptide in attenuating phyB sensing, signal transduction, or response.

This observation provided an excellent starting point to examine the precise mechanism of the peptide’s effect, because red light effects on plant development and gene expression are well described, both genetically and physiologically. Red light responses are typically mediated by phytochrome B (phyB). The receptor is activated, translocated, and accumulates in the nucleus within several hours of illumination (Yamaguchi et al., 1999; Gil et al., 2000), where it affects gene expression (Tepperman et al., 2004). Mechanistically, phyB directs various nuclear red light regulated responses through interaction with Phytochrome Interacting Factors (PIFs) (Ni et al., 1999; Huq and Quail, 2002; Kim et al., 2003; Huq et al., 2004). The light-activated phyB receptor binds to PIFs, in some cases preventing their association with promoter sequences in target genes (Huq and Quail, 2002) while controlling the proteolytic degradation of others (Bauer et al., 2004). The intricate coordination of signaling steps, re-localization of proteins, DNA binding, protein turnover, then integration into hormonal response, changes in turgor and cell wall plasticity, present many opportunities for interference from a rogue chemistry. Precise characterization of the localization and timing of PEP6-32’s effect on plant development would potentially provide a starting point for discovery of interactors, as well as a basis to further describe detailed facets of red light response. Therefore we tested the hypothesis that the synthetic peptide PEP6-32 interfered with some aspect of phyB-mediated signaling.

They hypothesis was tested by examining a set of well characterized phyB-mediated responses, and the results indicate that the PEP6-32 peptide is playing a role in discrete tissues that may or may not be directly connected to phyB response. Hypocotyl growth inhibition and cotyledon expansion were clearly impaired during red-light-mediated development, yet less so under blue or white light. These results support the hypothesis that the response is related to phyB.

However, examination of other classical phyB-mediated responses do not support the hypothesis. Wild type seedlings grown under red light on vertical plates exhibit directional growth abnormalities, as hypocotyls grow in various directions relative to the gravitational vector. However, *phyB* mutants grow straight up on vertical plates in red light (Poppe et al., 1996; Kim et al., 2011). Seedlings expressing PEP6-32 do not show a significantly different phenotype relative to non-transgenic controls (Figure 2), so while some facets of early red light signaling are impaired, others are completely unaffected. This finding indicates that the effect is not occurring at the level of the receptor or primary signaling events, or that the directional growth effect develops separately from stem growth inhibition and cotyledon expansion where the effect of the peptide is observed.

It is also well established that specific transcripts are induced by a short, single red light treatment in etiolated seedlings, under the direction of phyB (Kuno et al., 2000; Tepperman et al., 2004). Red light treatments induce steady-state transcript accumulation from members of the *Light-Harvesting, Chlorophyll-Binding* (*Lhcb*, also *cab*) gene family (Kaufman et al., 1984; Karlin-Neumann et al., 1988). The seedlings expressing PEP6-32 show normal transcript accumulation in response to red-light treatment. These results also indicate that PEP6-32 is not likely participating in red light signaling or integration, again not supporting the hypothesis of direct connections with phyB signal integration.

The effects on long-term phyB-mediated responses were also examined. Flowering is controlled by a variety of signals meant to assess plant age, season, local growing environment, and other factors that may impact seed production and viability (Putterill et al., 2004). In Arabidopsis, these signals primarily control flowering through the suppression or promotion of CONSTANS (CO) and FLOWERING LOCUS T (FT) expression and activity (Valverde et al., 2004). Red wavebands delay flowering by promoting CO degradation through a COP1-independent pathway mediated by phyB (Demotes-Mainard et al., 2016). Consequently, in *phyB* mutants, CO remains elevated, increasing expression of FT which can then induce flowering (Endo et al., 2005). Therefore, *phyB* mutant plants flower early. If the PEP6-32 had a role in phyB signalling later in development, seedlings would be expected to flower early, phenocopying *phyB* mutants. However, the plants expressing the peptide flowered at the same time or even slightly later than wild-type plants, which is exactly the opposite of what occurs in *phyB* mutants. The effect is minor yet statistically significant in some transgenic lines. The result is important in that it indicates that the effect of PEP6-32 in limiting red light response is likely limited to early seedling growth and is not exerting an impairment of all red-light-related responses.

Taken together the interpretation is that PEP6-32 is affecting cell expansion in the developing seedling, conditionally under red light. Hypocotyl growth rate inhibition and cotyledon expansion are repressed, two processes that are dependent on cell expansion, yet in opposing directions. Cotyledons typically expand upon illumination, yet here are smaller in the presence of the peptide after red light treatment. Hypocotyls are longer in the presence of the peptide when their elongation should be limited. It is tempting to speculate that the peptide could be functioning near the apical meristem, possibly by interacting with the hormone synthesis, transport or sensitivity that changes the location of expansion from the hypocotyl to cotyledon during the transition from darkness to light. The response is specific to red light, where the same expansion/inhibition patterns have been shown to be regulated in the same way by blue or far-red light. It is unclear why the effect is only observed under red light. One possibility is that blue and far-red signals excite alternative mechanisms that mask the red-light response.

The inhibition of red light seedling response was also induced by application of exogenous peptide. Seedlings grown on media containing the peptide exhibited longer hypocotyls. The effects were less pronounced in magnitude compared to transgenic seedlings expressing PEP6-32, but exhibited statistically-separable, dose-dependent action over two concentrations. These findings confirm that the phenotype is caused by the peptide and not another product of the transgene, such as an mRNA. Furthermore, this shows that the peptide can be taken up by seedlings to induce a phenotypic change.

The seminal work by Bao et al., (2017) illustrated that biologically active synthetic peptides may be identified by studying aberrancies in populations of random-DNA-expressing plants. This report expands these findings to characterize the effect of the PEP6-32 peptide, showing effects isolated to specific developmental windows, tissues and conditions, as well as effects induced by exogenous application. Future experiments will examine the structure-function aspects of the amino acid sequence to further probe biochemical activity and will test for specific protein-protein interactions. The trials presented in this report show that synthetic peptides created from random DNA sequence have the capacity to produce new molecules with discrete connections to plant biology. The results further validate this approach as a way to discover novel synthetic chemistries, as well as provide new tools to explore basic questions in plant growth, development and metabolism.

## Materials and Methods

### Plant Materials

The *PEP6-32* peptide expressing plants were grown from seeds generated previously as described in Bao et al., 2017. Corresponding non-transformed controls were obtained from segregating sibling seedlings not expressing a coincident GFP marker, or for some tests Col-0 was used as a non-transformed control. The phyB-5 mutant has been previously described (Reed et al., 1993).

### Hypocotyl elongation assay

Seeds were surface sterilized with consecutive washes of 70% ethanol for 3 min, 10% bleach for 15 min, then five rinses with sterile water. Seeds were then planted on minimal media (1 mM CaCl_2_, 1 mM KCl with 0.8% phytoagar) on square (100 mm × 100 mm) plates. Once planted, plates were stratified in darkness at 4°C for 4 d. The plates were then given 5 h of white light (~130 μmol⋅m^−2^⋅s^−1^) to induce germination and placed vertically in custom built LED light chambers under constant red light (30 μmol⋅m^−2^⋅s^−1^), constant blue light (10 μmol⋅m^−2^⋅s^−1^), or complete darkness for 96 h. Fluence rate measurements were taken using a Li-Cor LI-250 light meter. After the light treatment, the plates were scanned on a flatbed scanner and ImageJ was used to measure individual hypocotyls against an imaged standard. Relative hypocotyl length was calculated by dividing hypocotyl length of light grown plants by the hypocotyl length of dark grown plants.

### Seedling directional growth

IC Measure Software (The Imaging Source, LLC) was used to measure the plant angle of growth from images taken during 3 previous hypocotyl elongation assays. Multiple measurements were taken, and an average calculated for plants that switched major direction of growth mid development. The average angle of growth was calculated by converting each individual angle to 0-180° where 0° is vertical against the direction of gravity and 180° is straight down, in the direction of gravity.

### Cotyledon expansion assay

Seeds were surface sterilized by spraying them in 70% ethanol on paper and allowing to dry in a laminar airflow hood. Seeds from two independent transgenic lines were then plated on a deep Petri dishes (100 mm × 25 mm) containing 0.5x MS basal medium (cat. RPI #M10200) and 0.8% phytoagar with each line covering one half of the plate. Afterwards, the plates were stratified in darkness at 4°C for 4 d. The plates were then given 5 h of white light (~130 μmol⋅m^−2^⋅s^−1^) to induce germination and placed horizontally in light chambers at 96 μmol⋅m^−2^⋅s^−1^ white light, 30 μmol⋅m^−2^⋅s^−1^ red, or 10 μmol⋅m^−2^⋅s^−1^ blue under 16 h light conditions for 7 d. After treatment, seedlings were randomly chosen and cotyledons were removed under a dissecting microscope and imaged. ImageJ software was used to measure the area of each cotyledon. Between 72-83 cotyledons were analysed per genotype.

### Time to flower assay

Seeds were surface sterilized and planted on square plates containing 0.5x MS basal media, solidified with 0.8% phytoagar. The plates were stratified in darkness at 4°C for 4 d and placed vertically under white light (~130 μmol⋅m^−2^⋅s^−1^). At 7 d after germination, uniform seedlings were transferred to soil. In addition, the *PEP6-32* seedlings were screened to confirm GFP expression (verifying transgene presence) before planting. Each 10 cm pot containing soilless mix (Fafard 2P) contained two independent lines with three plants per line and each line was replicated in three pots. The plants were then grown in a growth chamber (Percival Model E36L) under long day (16 h day, 8 h night) conditions at a temperature of 21°C. Flowering was scored as when the inflorescence was 1 cm in length or greater.

Flowering time was also examined under enriched red light conditions. Ambient light spectra were measured in a fluorescent growth chamber using a StellarNet Inc spectroradiometer (Model EPP2000) and SpectraWiz Software. The conditions were approximated using LED – based illumination to best match the fluorescent light spectrum. Two additional light chambers were set to the same spectrum, and the amount of red (660nm) light was increased to 220% and 325% while decreasing 590nm light to maintain the same overall PAR (see supplemental Fig. 1). Plants were prepared using the same method as above except the plates were grown in the respective LED light chamber (control white, red 220%, or red 325%) prior to transplanting into pots.

### Gene expression

Red light induces transcripts from specific genes in etiolated seedlings rapidly after treatment. To measure the expression of the red light regulated genes, seeds were surface sterilized and planted on small Petri dishes (60mm × 15mm) containing 0.5 × MS media. The plates were stratified in darkness at 4°C for 4 d and placed under white light (~130 μmol⋅m^−2^⋅s^−1^) for 5 h to induce germination. Afterwards, the plates were placed in darkness at room temperature for four days. The etiolated seedlings were then exposed to 8 μmol⋅m^−2^⋅s^−1^ of red light for 1 h in the LED light chamber or placed into complete darkness. Following treatment, plants were immediately frozen in liquid nitrogen. The shoots were ground to a fine powder with cold mortar and pestle and RNA was isolated using a Qiagen RNeasy kit. The RNA was treated with Promega RQ1 DNase and reverse transcribed into cDNA using the Improm-II Reverse Transcription kit (Promega; Fitchberg, WI). Steady-state transcript levels were then measured using the cDNA template in quantitative PCR (qPCR) reactions with SYBR Green reagents. Ubiquitin Family Protein (*UFP*) was used as the endogenous control. Supplemental Table 1 shows sequences of *CAB2* and *LHCB1.5* primers used. The fold change in gene expression reported is an average of three RT-qPCR replicates.

For peptide expression analysis, mature leaves of plants were harvested and flash frozen. Next, 0.1 g of tissue was used in the protocol described above. Peptide expression was then compared relative to Line 1. The expression levels reported are an average of two RT-qPCR runs.

### Exogenous application of synthesized peptide

The PEP6-32 peptide (MACPASVSVC) was synthesized commercially and dissolved in a buffer containing 1 mM CaCl_2_, 1 mM KCl, and 10 mM HEPES. The peptide concentration was determined with a spectrophotometer. Seeds were surface sterilized and stored in 1.5 mL Eppendorf tubes with 1 mL of sterile water for stratification in darkness at 4°C for 3 d. Afterwards, the seeds were planted on square plates containing minimal media with added peptide at a final concentration of 1 μM or on control plates with an equivalent amount of buffer without peptide. The plates were then placed under white light (~130 μmol⋅m^−2^⋅s^−1^) for 5 h. After which, they were transferred to LED light chambers set to 1, 5, 10, 30, or 100 μmol⋅m^−2^⋅s^−1^ of constant red light for 96 h or kept dark as a control. Hypocotyl lengths were measured as described above.

### Statistical Analysis

The Mann-Whitney U Test (two-tailed) was used for all statistical analysis.

## Author Contributions

T.S. Performed all experiments that produced data for this report, and provided the first draft of the manuscript; R.C. identified and isolated the original plant line studied; Z.B. participated in generation of plant materials and supervision of the experiments; M.C. designed and prepared the constructs for transformation and developed the random peptide-expressing lines; K.F. conceived the project and wrote the article with contributions from all of the authors, and agrees to serve as the author responsible for contact and ensures communication.

## Acknowledgements

(none)

---

**Supplemental Figure 1. Light spectrum of the different growth areas.** Light spectrum of the growth chamber (A), LED light chamber set to approximate spectrum of growth chamber (B), LED light chamber enriched for 220% red (C), LED light chamber enriched for 325% red (D), individual spectrum of each configurable LED setting when set to 10% power (E), table indicating the fluence rate of each wavelength (F). The LED light chambers can be configured for variable power (0-100%) for 470nm, 530nm, 590nm, 660nm, and 730nm. (E) shows the spectrum given off when one of the wavelengths is set to 10% and the rest to 0%. This information was used to determine the wavelength bins to measure in the growth chamber for matching in the LED light chambers. (C) and (D) increased the amount of 660nm (bin 630-678nm) at the cost of 590nm (bin 565-639nm) to maintain the total PAR.

**Supplemental Table 1.** Sequences of primers used in RT-qPCR

## Literature Cited

Bao Z, Clancy MA, Carvalho RF, Elliott K, Folta KM (2017) Identification of Novel Growth Regulators in Plant Populations Expressing Random Peptides. Plant Physiol 175: 619–627

Bauer D, Viczián A, Kircher S, Nobis T, Nitschke R, Kunkel T, Panigrahi KCS, Ádám É, Fejes E, Schäfer E, Nagy F (2004) Constitutive Photomorphogenesis 1 and Multiple Photoreceptors Control Degradation of Phytochrome Interacting Factor 3, a Transcription Factor Required for Light Signaling in Arabidopsis. The Plant Cell 16: 1433

Chen M, Chory J (2012) Phytochrome signaling mechanisms and the control of plant development. Trends in Cell Biology 21: 664–671

De Rybel B, Audenaert D, Vert G, Rozhon W, Mayerhofer J, Peelman F, Coutuer S, Denayer T, Jansen L, Nguyen L, Vanhoutte I, Beemster GTS, Vleminckx K, Jonak C, Chory J, Inzé D, Russinova E, Beeckman T (2009) Chemical Inhibition of a Subset of Arabidopsis thaliana GSK3-like Kinases Activates Brassinosteroid Signaling. Chemistry & Biology 16: 594–604

Demotes-Mainard S, Péron T, Corot A, Bertheloot J, Le Gourrierec J, Pelleschi-Travier S, Crespel L, Morel P, Huché-Thélier L, Boumaza R (2016) Plant responses to red and far-red lights, applications in horticulture. Environmental and Experimental Botany 121: 4–21

Endo M, Nakamura S, Araki T, Mochizuki N, Nagatani A (2005) Phytochrome B in the mesophyll delays flowering by suppressing FLOWERING LOCUS T expression in Arabidopsis vascular bundles. The Plant Cell 17: 1941–1952

Fiers M, Hoogenboom J, Brunazzi A, Wennekes T, Angenent GC, Immink RGH (2017) A plant-based chemical genomics screen for the identification of flowering inducers. Plant Methods 13: 78

Gil P, Kircher S, Adam E, Bury E, Kozma-Bognar L, Schafer E, Nagy F (2000) Photocontrol of subcellular partitioning of phytochrome-B:GFP fusion protein in tobacco seedlings. Plant J 22: 135–145

Hagihara S, Yamada R, Itami K, Torii KU (2019) Dissecting plant hormone signaling with synthetic molecules: perspective from the chemists. Current Opinion in Plant Biology 47: 32–37

Huq E, Al-Sady B, Hudson M, Kim C, Apel K, Quail PH (2004) PHYTOCHROME-INTERACTING FACTOR 1 is a critical bHLH regulator of chlorophyll biosynthesis. Science 305: 1937–1941

Huq E, Quail PH (2002) PIF4, a phytochrome-interacting bHLH factor, functions as a negative regulator of phytochrome B signaling in Arabidopsis. Embo J 21: 2441–2450

Karlin-Neumann GA, Sun L, Tobin EM (1988) Expression of Light-Harvesting Chlorophyll a/B-Protein Genes Is Phytochrome-Regulated in Etiolated Arabidopsis-Thaliana Seedlings. Plant Physiology 88: 1323–1331

Kaufman LS, Thompson WF, Briggs WR (1984) Different Red Light Requirements for Phytochrome-Induced Accumulation of cab RNA and rbcS RNA. Science 226: 1447–1449

Kim J, Yi H, Choi G, Shin B, Song P-S, Choi G (2003) Functional Characterization of Phytochrome Interacting Factor 3 in Phytochrome-Mediated Light Signal Transduction. The Plant Cell 15: 2399

Kim K, Shin J, Lee S-H, Kweon H-S, Maloof JN, Choi G (2011) Phytochromes inhibit hypocotyl negative gravitropism by regulating the development of endodermal amyloplasts through phytochrome-interacting factors. Proceedings of the National Academy of Sciences 108: 1729–1734

Kuno N, Muramatsu T, Hamazato F, Furuya M (2000) Identification by large-scale screening of phytochrome-regulated genes in etiolated seedlings of Arabidopsis using a fluorescent differential display technique. Plant Physiol 122: 15–24

Mayfield JD, Folta KM, Paul AL, Ferl RJ (2007) The 14-3-3 Proteins mu and upsilon influence transition to flowering and early phytochrome response. Plant Physiol 145: 1692–1702

Ni M, Tepperman JM, Quail PH (1999) Binding of phytochrome B to its nuclear signalling partner PIF3 is reversibly induced by light. Nature 400: 781–784

Parks BM, Spalding EP (1999) Sequential and coordinated action of phytochromes A and B during Arabidopsis stem growth revealed by kinetic analysis. Proc Natl Acad Sci U S A 96: 14142–14146

Poppe C, Hangarter RP, Sharrock RA, Nagy F, Schafer E (1996) The light-induced reduction of the gravitropic growth-orientation of seedlings of Arabidopsis thaliana (L.) Heynh. is a photomorphogenic response mediated synergistically by the far-red-absorbing forms of phytochromes A and B. Planta 199: 511–514

Quail PH (2002) Phytochrome photosensory signalling networks. Nat Rev Mol Cell Biol 3: 85–93

Reed JW, Nagpal P, Poole DS, Furuya M, Chory J (1993) Mutations in the gene for the red/far-red light receptor phytochrome B alter cell elongation and physiological responses throughout Arabidopsis development. Plant Cell 5: 147–157

Robson P, Whitelam GC, Smith H (1993) Selected Components of the Shade-Avoidance Syndrome Are Displayed in a Normal Manner in Mutants of Arabidopsis thaliana and Brassica rapa Deficient in Phytochrome B. Plant Physiol 102: 1179–1184

Somers DE, Sharrock RA, Tepperman JM, Quail PH (1991) The hy3 Long Hypocotyl Mutant of Arabidopsis Is Deficient in Phytochrome B. Plant Cell 3: 1263–1274

Tepperman JM, Hudson ME, Khanna R, Zhu T, Chang SH, Wang X, Quail PH (2004) Expression profiling of phyB mutant demonstrates substantial contribution of other phytochromes to red-light-regulated gene expression during seedling de-etiolation. Plant J 38: 725–739

Valverde F, Mouradov A, Soppe W, Ravenscroft D, Samach A, Coupland G (2004) Photoreceptor regulation of CONSTANS protein in photoperiodic flowering. Science 303: 1003–1006

Whitelam GC, Patel S, Devlin PF (1998) Phytochromes and photomorphogenesis in Arabidopsis. Philos Trans R Soc Lond B Biol Sci 353: 1445–1453

Whitelam GC, Smith H (1991) Retention of phytochrome-mediated shade avoidance responses in phytochrome-deficient mutants of Arabidopsis, cucumber and tomato. Journal of Plant Physiology 139: 119–125

Xu X, Paik I, Zhu L, Huq E (2015) Illuminating Progress in Phytochrome-Mediated Light Signaling Pathways. Trends in Plant Science 20: 641–650

Yamaguchi R, Nakamura M, Mochizuki N, Kay SA, Nagatani A (1999) Light-dependent translocation of a phytochrome B-GFP fusion protein to the nucleus in transgenic Arabidopsis. J Cell Biol 145: 437–445

Zhang C, Brown MQ, van de Ven W, Zhang Z-M, Wu B, Young MC, Synek L, Borchardt D, Harrison R, Pan S, Luo N, Huang Y-mM, Ghang Y-J, Ung N, Li R, Isley J, Morikis D, Song J, Guo W, Hooley RJ, Chang C-eA, Yang Z, Zarsky V, Muday GK, Hicks GR, Raikhel NV (2016) Endosidin2 targets conserved exocyst complex subunit EXO70 to inhibit exocytosis. Proceedings of the National Academy of Sciences 113: E41

