## Supplementary figures and images for "A Synthetic Peptide Encoded by a Random DNA Sequence Inhibits Discrete Red Light Responses"

### Supplemental Figure 1

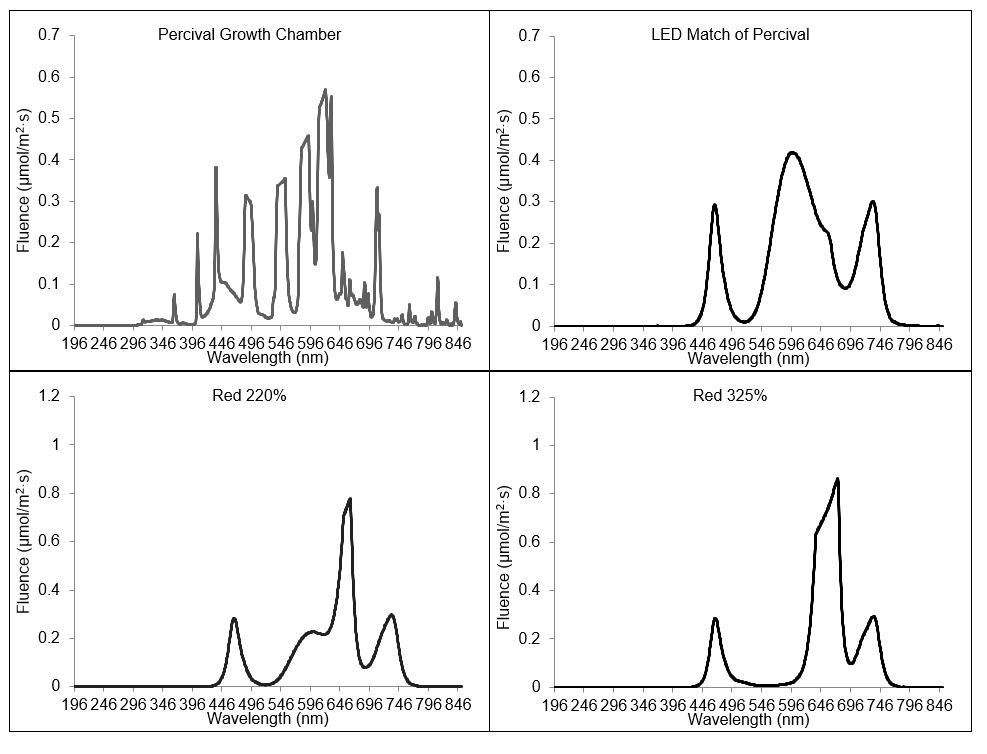

### Supplemental Table 2

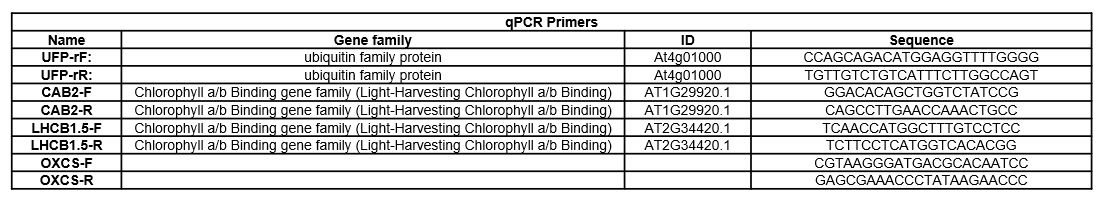
